# Lactate cannot replace glucose for maintaining the viability of mouse and human glioma cells

**DOI:** 10.1101/2025.06.03.655413

**Authors:** Eric Y. Miller, Tomás Duraj, Nathan L. Ta, Derek C. Lee, Purna Mukherjee, Oscar Shaver, Thomas N. Seyfried

## Abstract

**Objectives:** Aerobic lactic acid fermentation (the “Warburg effect”) is associated with OxPhos insufficiency and altered energy metabolism in most cancers. Whether lactate is a major fuel in cancer cells remains debated. This study investigated whether lactate could serve as a metabolic fuel in glioma cells and replace glucose to support viability.

**Methods:** A bioluminescence ATP assay and calcein-AM/EthD-III double-staining were used to measure ATP content and viability in mouse (VM-M3, CT-2A) and human (U-87MG) glioma cells differing in cell biology and genetic background. Viability was assessed in thioglycollate-elicited peritoneal macrophages (TPMs) from VM/Dk and C57BL/6J mice, used as syngeneic non-neoplastic controls for VM-M3 and CT-2A gliomas, respectively. Oxygen consumption rate (OCR) was determined using the Resipher system.

**Results:** Lactate alone failed to sustain ATP content and viability in all glioma cell lines. ATP content and viability were lower in cancer cells cultured in glutamine and lactate than in glucose and glutamine. In contrast, lactate alone sustained over 45% viability in both TPM models. Moreover, viability was similar between TPMs cultured in glutamine and lactate and in glucose and glutamine. In human U-87MG, lactate addition under severe glucose restriction increased OCR and viability. A low dose of the glycolysis inhibitor 2-deoxy-D-glucose abolished both increases.

**Conclusions:** Our results suggest non-neoplastic mouse TPMs utilize lactate more effectively than mouse glioma cells. In U-87MG, lactate utilization appears glycolysis-dependent, given its sensitivity to 2-deoxy-D-glucose. In conclusion, our data does not support lactate as a major oxidative fuel for viability in mouse and human glioma cells.

## Introduction

Lactate, traditionally regarded as a waste product of glucose fermentation, has gained significant attention in cancer metabolism due to its extracellular accumulation linked to aerobic lactic acid fermentation, or the “Warburg effect” [1–4]. The Warburg effect involves the reduction of pyruvate to lactic acid even in the presence of sufficient oxygen (“aerobic”), which contrasts with the complete oxidation of pyruvate to CO_2_ and water via oxidative phosphorylation (OxPhos) observed in most non-tumoral cells. Defects in mitochondrial number, structure, or function have been observed in all major cancers, suggesting a linkage between mitochondrial dysfunction, aerobic fermentation, and neoplasia [5,6]. Mitochondrial dysfunction would be expected to impair OxPhos, and therefore lactate oxidation, based on the biological principle that structure determines function [5,7,8]. For instance, the number and structure of cristae are closely linked to the efficiency of the electron transport chain, and partial or total cristolysis has been documented in all major cancers [8–13]. Alternative hypotheses suggest that the Warburg effect might arise as an adaptation to increased redox stress or biosynthetic demands [14]. Hypotheses based on teleology, or design with purpose, suggest that tumor cells choose aerobic fermentation over OxPhos and that the Warburg effect provides an overall benefit by supporting a tumor microenvironment (TME) conducive to cancer cell proliferation [15,16]. Regardless of purpose, the Warburg effect is a common phenotype of all major cancers [5,14,17–19].

Recent studies in glioblastoma (GBM) and other cancers have found that carbon-labeled lactate could contribute to the TCA cycle, with some models suggesting that the contribution of lactate might exceed that of glucose [20–23]. Other reports, however, indicate that glucose remains the primary contributor after accounting for isotopic exchange between pyruvate and lactate [23]. Cai et al. proposed that lactate might act as a mitochondrial signaling molecule, stimulating OxPhos in cancer cells independent of its metabolism [24]. Torrini et al. recently suggested that lactate could be an important oxidative fuel under the assumption that OxPhos is unimpaired in GBM cells [25]. The Lisanti group also suggested that cancer-associated fibroblasts secrete lactate, which is then taken up and oxidized by cancer cells, in what has been coined the “Reverse Warburg Effect” [26,27].

Regardless of mechanism, substantial oxidation of lactate for ATP production would be unexpected in neoplastic cancer cells given the rapid export of lactate via the Warburg effect [2,28–30], as well as the aforementioned mitochondrial defects found in all major cancers [5,12,31]. Elevated lactic acid levels are a consistent feature of the TME, with extracellular lactic acid concentrations reaching up to 40 mM, approximately 20 times higher than the levels found in blood [32,33]. Although elevated lactic acid levels are associated with poor clinical outcomes in some cancers [34–36], targeting lactate oxidation as a cancer treatment risks conflating cause and effect. Likewise, while lactate carbons could enter the TCA cycle, the relative contribution of lactate to mitochondrial metabolism in cancer remains controversial [22,23]. Furthermore, lactate-driven anaplerosis could occur without a corresponding increase in ATP production via OxPhos, as TCA cycle turnover is not always coupled to OxPhos in cancer cells [37,38]. Thus, it is important to characterize the functional impact of lactate on the survival of glioma cells by assessing viability and ATP production under different nutrient conditions.

In this study, we used a luciferin-luciferase bioluminescence ATP assay and a calcein-AM/EthD-III double-staining assay to evaluate the effects of lactate supplementation on the ATP content and viability of mouse and human glioma cell lines. We also used the calcein-AM/EthD-III double-staining assay to evaluate the viability of primary non-neoplastic TPMs. Notably, VM-M3 glioma cells and VM/Dk TPMs share both a common genetic background (mouse strain) and myeloid cell biology (mesenchymal origin) [39], while CT-2A glioma cells and C57BL/6J TPMs share a common genetic background but differ in cell biology (stem cell glioma versus macrophage, respectively) [40,41]. We found that metabolic flexibility in lactate utilization was greater in the C57BL/6J and VM/Dk TPMs than in the mouse (CT-2A and VM-M3) or human (U-87MG) glioma cells. In contrast to the non-neoplastic mouse TPMs, where lactate could substitute for glucose to support viability in the presence of glutamine, lactate was unable to replace glucose to support viability of the mouse and human glioma cells or maintain viability in the absence of glucose and glutamine. These results highlight the metabolic differences between neoplastic glioma cells and non-neoplastic TPMs.

## Methods

### Mice

Mice of the VM/Dk (VM) inbred strain were originally obtained as gifts from G. Carlson (McLaughlin Research Institute, Great Falls, Montana) and from H. Fraser (University of Edinburgh, Scotland). Mice of the C57BL/6J (B6) inbred strain were obtained originally from the Jackson Laboratory, Bar Harbor, ME. All mice used in this study were housed and bred in the Boston College Animal Care Facility under IACUC-approved husbandry conditions as previously described [42]. Male and female mice between 8–10 weeks of age were used for all studies. All animal procedures and protocols were in strict accordance with the NIH Guide for the Care and Use of Laboratory Animals and approved by the Institutional Animal Care Committee at Boston College under assurance number A3905–01.

### Thioglycollate-elicited peritoneal macrophages

The protocol for generating and collecting TPMs was adapted from a previously described method [43]. In brief, 8–10-week-old mice were intraperitoneally injected with 1 mL of sterile 3% thioglycollate broth (Sigma-Aldrich, St. Louis, MO). After 96 hours, mice were euthanized, and the peritoneal cavity was lavaged with 10 mL of DMEM. The lavage fluid was collected and centrifuged at 300 × g for 5 minutes, and the cell pellet was resuspended in seeding medium for 2 hours. Initial seeding medium was Dulbecco’s Modified Eagle Medium (DMEM) with sodium bicarbonate, without glucose, glutamine, or serum (D5030; Sigma-Aldrich, St. Louis, MO), supplemented with 2% penicillin-streptomycin (15140-122; Gibco, Waltham, MA). Non-adherent cells and red blood cells were removed by tapping and washing three times with PBS, leaving an enriched macrophage population for subsequent analyses.

### Cell lines

The VM-M3 cell line was established from a spontaneously arising tumor in the cerebrum of an adult VM/Dk inbred mouse as previously described [39]. The VM-M3 cell line exhibits all of the growth characteristics typical of human high-grade gliomas, including invasion patterns known as the secondary structures of Scherer [44]. The CT-2A cell line was established from a tumor induced via 20-methylcholanthrene in the cerebral cortex of a C57BL/6J mouse [40]. CT-2A was initially classified as a malignant anaplastic astrocytoma, while more recent studies suggest similarities to neoplastic neural stem cells [41]. The U-87MG cell line was obtained as a gift from Miguel Sena-Esteves (UMass Medical Center, Worcester, MA). The U-87MG cell line is a well-established human glioblastoma cell model [45–47]. All cell lines were transfected with a lentivirus vector containing the firefly luciferase gene under the control of the cytomegalovirus promoter as previously described, resulting in the cell lines CT-2A/Fluc, VM-M3/Fluc, and U-87MG/Fluc [39,48]. All cell lines were routinely tested and confirmed to be free of mycoplasma.

### Culture conditions

Tumor cell stock plates were maintained in seeding medium. Cells were maintained in a humidified incubator at 37°C and 5% CO_2_. Experimental media was made from glucose-free and glutamine-free powdered DMEM (D5030; Sigma-Aldrich), supplemented with nutrients corresponding to each experimental condition. All experimental media was supplemented with 10% fetal bovine serum (FS-0500-AD, lot: S12E22AD1; Atlas Biologicals, Fort Collins, CO). For experimental use, tumor cells were seeded in seeding medium and allowed to settle for 24 hours. The seeding medium was then replaced by the corresponding experimental media. Cells were incubated in the experimental media at 37°C and 5% CO_2_ for the duration specified in each experimental protocol.

### Luciferin-luciferase bioluminescence assay

Following the designated incubation time in experimental media, luciferin (20 mg/mL, Syd Labs, Hopkinton, MA) was added to each well containing the luciferase-tagged cells and allowed to incubate at room temperature for 1 minute, as previously described [39]. Plates were read with the Ami HT (Spectral Instruments, Tucson, AZ) imaging device after a two-minute exposure. Bioluminescence was quantified using the Aura software (version 2.3.1, Spectral Instruments, Tucson, AZ). Photon values were obtained in Aura by selecting a region of interest for each well following manufacturer instructions.

### Ethidium homodimer III and calcein-AM assay

Ethidium homodimer III (EthD-III) and calcein-AM were used to visualize cell viability [49]. EthD-III is a membrane-impermeant dye that emits red fluorescence when it binds to nucleic acids. As EthD-III only enters cells when the plasma membrane is compromised, it marks dead or late apoptotic cells. Calcein-AM is a membrane-permeable dye that is hydrolyzed by intracellular esterases in live cells to produce green fluorescence. Cells exhibiting only red fluorescence are non-viable, whereas cells emitting only green fluorescence are viable. Cells fluoresce both colors where membrane permeabilization has begun but esterase activity still remains [49]. This indicates cells that are apoptotic, or cells undergoing loss of viability such that esterase activity has not yet ceased [50,51]. Tumor cells were seeded, transitioned to experimental media, and incubated in the experimental media for the designated period as outlined in *Culture conditions*. After incubation, a freshly prepared 50 µL DMEM-based solution containing 2.0 µM EthD-III and 1.0 µM calcein-AM was added to each well. Plates were incubated for 45 minutes at 37°C. Fluorescence images were acquired using the EVOS FL Imaging System (Life Technologies, Carlsbad, CA) with a 10× objective and Texas Red or GFP filter cubes. In some cases, CellProfiler software was used to algorithmically quantify the live vs. dead ratio.

### Oxygen consumption rate assay

Oxygen consumption rate (OCR) was measured continuously using the Resipher instrument (Lucid Scientific, Atlanta, GA) in a 96-well flat-bottom plate. Cells were seeded at a density of 2.0 x 10^5^ cells/well and changed to experimental media after 24 hours. For calibration, the sensing lid of the Resipher was placed on top of the plate for five minutes prior to starting OCR measurement. OCR measurement was started in Lucid Lab software (Lucid Scientific, Atlanta, GA) with the sensing lid remaining on the plate. Cells were maintained during measurement in a humidified incubator at 5% CO_2_ and 90-95% relative humidity. OCR was monitored for 72 hours. Data was analyzed using Lucid Lab software.

### Statistics

Statistical analyses and data visualization were performed using Prism (version 9; GraphPad, La Jolla, CA). Data was analyzed using one-way analysis of variance (ANOVA), with Tukey’s correction for multiple comparisons. In all figures, error bars represent mean ± standard deviation (SD), and *n* indicates the number of biological replicates. A *p* ≤ 0.05 was considered statistically significant. Significance levels are denoted as ∗ *p* ≤ 0.05, ∗∗ *p* ≤ 0.01, ∗∗∗ *p* ≤ 0.001, and ∗∗∗∗ *p* ≤ 0.0001.

## Results

The aim of this research was to determine whether lactate could serve as a fuel source supporting ATP content and viability in glioma cells. A luciferin-luciferase bioluminescence ATP assay was used to evaluate the effects of glucose, glutamine, and lactate on ATP content and viability in mouse (VM-M3, CT-2A) and human (U-87MG) glioma cells. This assay has been validated as a reliable marker of viable cell number over time, demonstrating a strong correlation between bioluminescence emission and the number of viable cells [52]. In viable luciferase-tagged cells, luciferase catalyzes the reaction between luciferin and ATP, producing oxyluciferin and light, with luminescence intensity proportional to ATP levels in the sample [53].

### Lactate fails to substitute for glucose in the VM-M3 glioma cells but supports viability in VM/Dk TPMs

To compare the effects of lactate on the viability of the VM-M3 glioma cells with that of a non-neoplastic control, we evaluated the effects of lactate-containing media on the viability of both the VM-M3 glioma cells and the non-neoplastic VM/Dk-derived TPMs. The cells were cultured for 48 hours in DMEM alone (negative control), glucose and glutamine (positive control), lactate alone, glutamine alone, or glutamine and lactate (Figure 1A). Bioluminescence of the VM-M3 cells cultured with lactate alone was negligible and indistinguishable from that of cells in DMEM alone. Bioluminescence was lower in the VM-M3 cells cultured in glutamine and lactate than in the cells cultured in glucose and glutamine. To further validate the bioluminescence assay as a marker of cell viability, we also quantified viability using calcein-AM and EthD-III fluorescent staining of the VM-M3 cells, with similar results (Figure S1).

**Figure 1.**
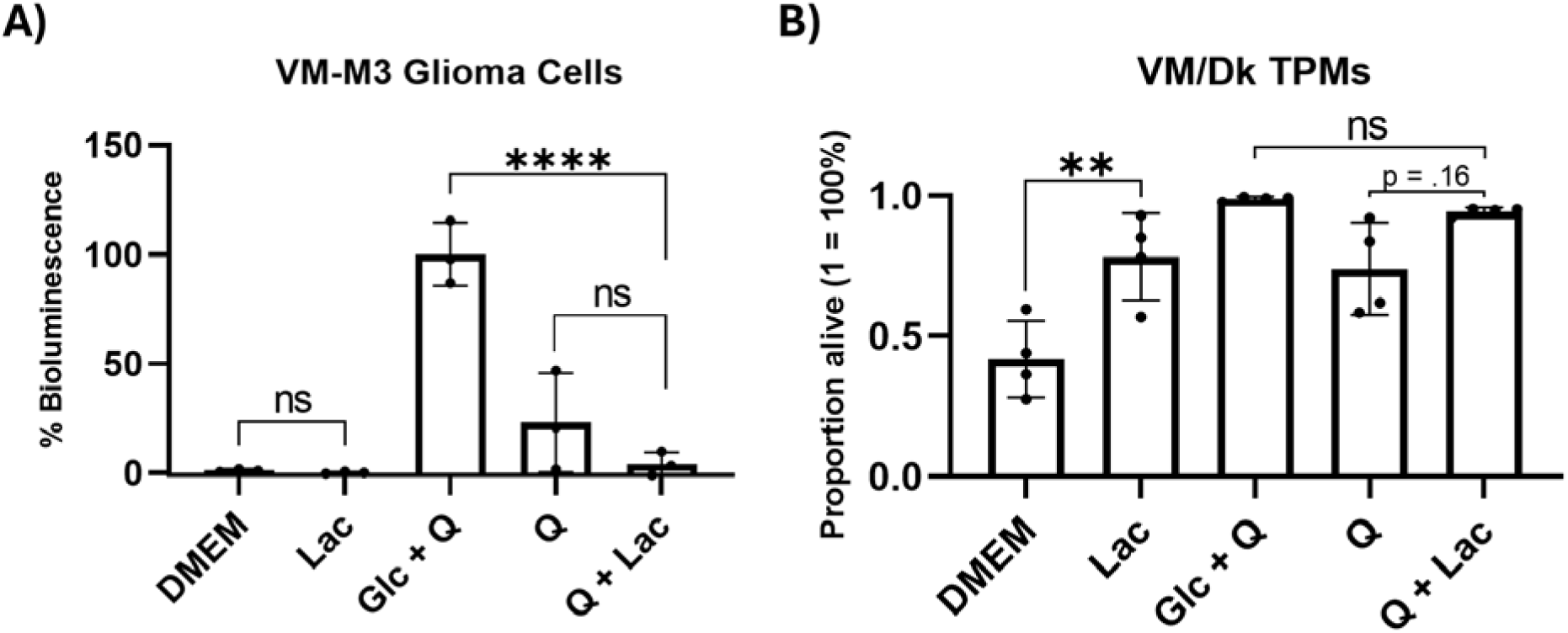
Effect of lactate on the viability of VM-M3 glioma cells and VM/Dk TPMs. (A) VM-M3 cells were seeded and cultured for 48 hours in the specified media. Bioluminescence was measured and normalized to the positive control (Glc + Q). (B) VM/Dk TPMs were seeded and cultured for 48 hours in the same media used for VM-M3 cells. Viability was quantified via CellProfiler using calcein-AM and EthD-III staining images. The media conditions were DMEM (negative control), glucose and glutamine (Glc + Q; positive control), lactate (Lac), glutamine (Q), or glutamine and lactate (Q + Lac). When present, glucose, lactate, and glutamine concentrations were 12.0 mM, 12.0 mM, and 2.0 mM, respectively. Error bars represent mean ± SD with a minimum of 3 independent experiments. Significance levels are denoted as ∗ *p* ≤ 0.05, ∗∗ *p* ≤ 0.01, ∗∗∗ *p* ≤ 0.001, and ∗∗∗∗ *p* ≤ 0.0001. These results show that lactate could not replace glucose for maintaining the viability of VM-M3 glioma cells but could support the viability of the syngeneic, non-neoplastic VM/Dk TPMs.

In contrast to the VM-M3 glioma cells, viability was 36% greater in the VM/Dk TPMs cultured in lactate alone than when cultured in DMEM alone (Figure 1B). The VM/Dk TPMs cultured in glutamine and lactate did not exhibit greater viability than TPMs cultured in glutamine alone (*p* = 0.16). Viability in glutamine and lactate approached 100%, suggesting that additional increases might have been limited. The VM/Dk TPMs cultured in glutamine and lactate exhibited similar viability to TPMs cultured in glucose and glutamine. The calcein-AM and EthD-III fluorescent double-staining images illustrated the contrast between the VM-M3 glioma cells and the VM/Dk TPMs, showing near-complete cell death of VM-M3 cells cultured in lactate alone, whereas an average of 78% of VM/Dk TPMs survived in lactate alone (Figure S2). Taken together, these findings indicate that lactate alone could not sustain VM-M3 glioma viability over an extended period, nor could lactate replace glucose to sustain VM-M3 glioma viability. In contrast, lactate alone was able to sustain viability in VM/Dk TPMs.

### Lactate fails to substitute for glucose in the CT-2A glioma cells but supports viability in C57BL/6J TPMs

To compare the effects of lactate on the viability of the CT-2A glioma cells with that of a non-neoplastic control, we evaluated the effects of lactate-containing media on the viability of both the CT-2A glioma cells and the non-neoplastic C57BL/6J-derived TPMs, cultured under the same conditions described in Figure 1. The CT-2A cells exhibited results similar to those seen in the VM-M3 cells, as the addition of lactate did not increase bioluminescence under any condition (Figure 2A). Bioluminescence of the CT-2A cells cultured with lactate alone was negligible and indistinguishable from that of cells in DMEM alone. Bioluminescence was lower in the CT-2A cells cultured in glutamine and lactate than in the cells cultured in glucose and glutamine. Taken together with the results from the VM-M3 cells, these findings indicate that lactate is incapable of maintaining viability in two mouse glioma cell lines that differ in cell biology and genetic background.

**Figure 2.**
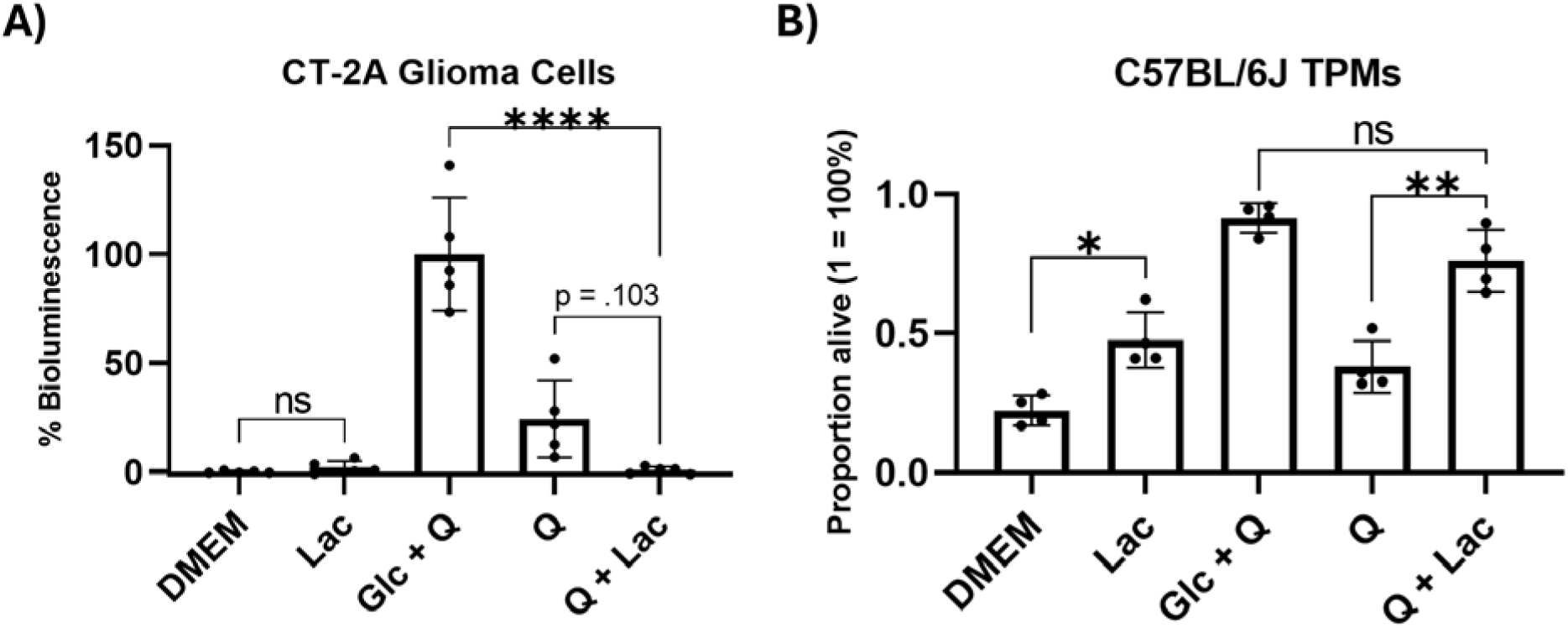
Effect of lactate on the viability of CT-2A glioma cells and C57BL/6J TPMs. (A) CT-2A cells were seeded and cultured for 48 hours in the specified media. Bioluminescence was measured and normalized to the positive control (Glc + Q). (B) C57BL/6J TPMs were seeded and cultured for 48 hours in the same media used for the CT-2A cells. Viability was quantified as in Figure 1. The media conditions were DMEM (negative control), glucose and glutamine (positive control; Glc + Q), lactate (Lac), glutamine (Q), or glutamine and lactate (Q + Lac). Other conditions, including statistics, are as described in Figure 1. These results show that lactate could not replace glucose for maintaining the viability of CT-2A glioma cells but could support the viability of the syngeneic, non-neoplastic C57BL/6J TPMs.

In contrast to the CT-2A glioma cells, viability was 25% greater in the C57BL/6J TPMs cultured in lactate alone than when cultured in DMEM alone (Figure 2B). The C57BL/6J TPMs cultured in glutamine and lactate exhibited a 38% absolute increase in viability compared to TPMs cultured in glutamine alone. The C57BL/6J TPMs cultured in glutamine and lactate exhibited similar viability to TPMs cultured in glucose and glutamine. The calcein-AM and EthD-III fluorescent double-staining images illustrated the contrast between the CT-2A glioma cells and the C57BL/6J TPMs, showing near-complete cell death of CT-2A cells cultured in lactate alone, whereas an average of 48% of C57BL/6J TPMs survived in lactate alone (Figure S3). Taken together, these findings indicate that lactate alone could not sustain CT-2A glioma viability over an extended period, nor could lactate replace glucose to sustain CT-2A glioma viability. In contrast, the addition of lactate to media containing C57BL/6J TPMs enhanced viability.

### Lactate fails to substitute for glucose in the human U-87MG glioma cells

We next evaluated the effects of lactate-containing media on the viability of the human glioma cell line U-87MG. The cells were cultured for 72 hours in DMEM alone (negative control), glucose and glutamine (positive control), lactate alone, glutamine alone, or in glutamine and lactate (Figure 3). Bioluminescence of the U-87MG cells cultured with lactate alone was negligible and indistinguishable from that of cells in DMEM alone. Bioluminescence was lower in the U-87MG cells cultured in glutamine and lactate than in the cells cultured in glucose and glutamine. The calcein-AM and EthD-III fluorescent staining images corroborated these patterns, showing complete cell death among U-87MG cells cultured in lactate alone (Figure S4). These findings indicate that lactate alone could not sustain viability in the human U-87MG glioma cells, nor could lactate replace glucose to sustain U-87MG glioma viability, consistent with the results found in the mouse CT-2A and VM-M3 glioma cells.

**Figure 3.**
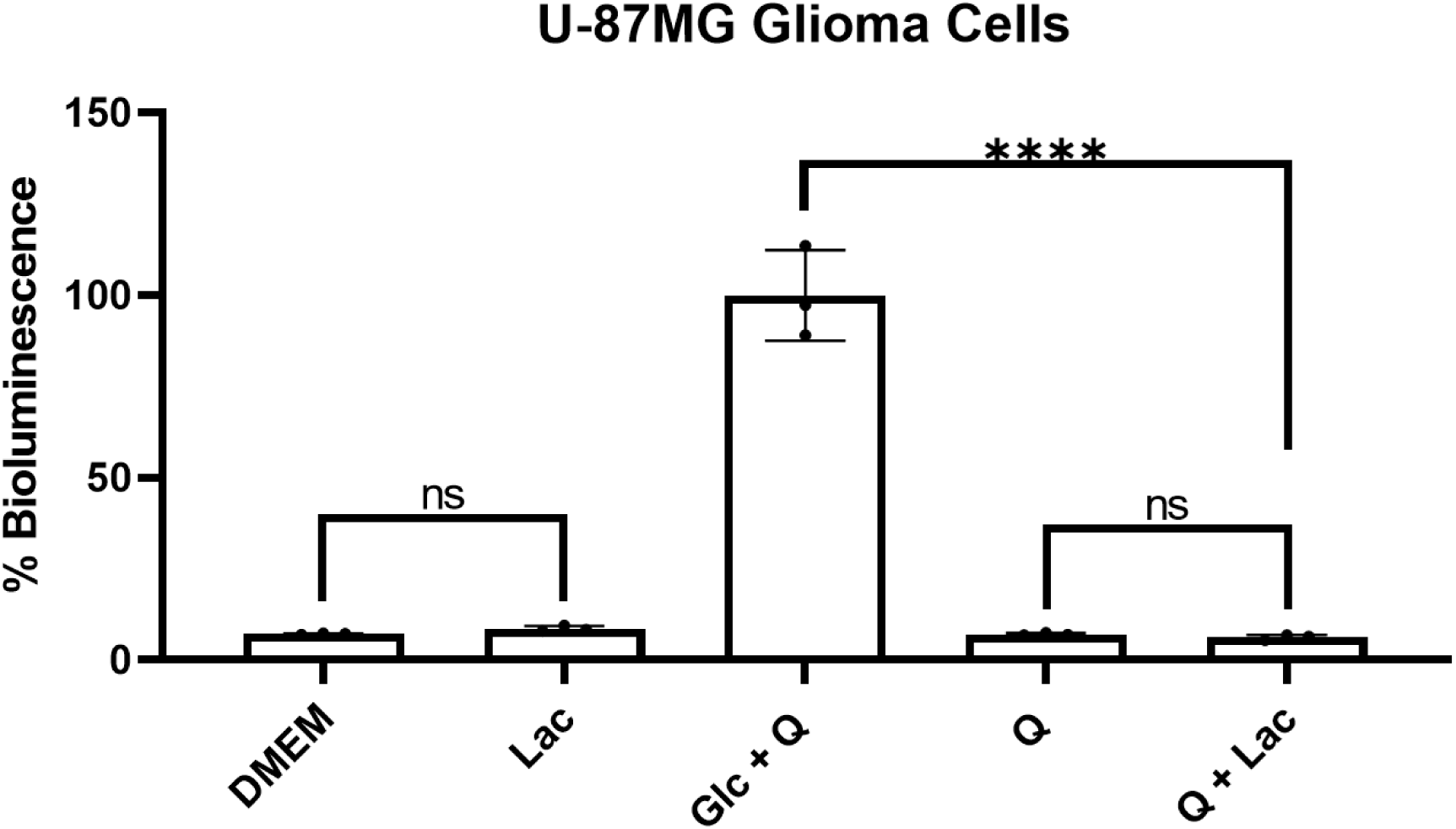
Effect of lactate on the ATP content and viability of U-87MG glioma cells. U-87MG glioma cells were seeded and cultured for 72 hours in the specified media. All quantifications were normalized to the positive control (12.0 mM glucose and 2.0 mM glutamine). To compare the effects of different concentrations of supplementary lactate, we tested a range of 5-15 mM lactate as a supplement to glutamine, along with testing 5-15 mM lactate as a supplement to glucose and glutamine (Figure S5). Supplementary lactate had no effect regardless of concentration. The concentration shown in “Q + Lac” is 15.0 mM lactate. Other conditions, including statistics, are as described in Figure 1. Taken together, these findings indicate that lactate alone could not sustain U-87MG glioma viability over an extended period, regardless of lactate concentration.

### Lactate supports viability under glucose restriction in U-87MG but not VM-M3 or CT-2A cells

In contrast to our findings with the VM-M3, CT-2A, and U-87MG glioma cells, previous research found that lactate could promote viability in neoplastic HepG2 and U-87MG cells [24,25]. At least two experimental factors might explain the differences between these findings: (1) media changes at regular intervals, or (2) low (but non-zero) glucose and/or glutamine in the lactate-containing media. Both situations involve constant or intermittent restriction of glucose and/or glutamine, creating periods where the availability of these fuels is insufficient to meet bioenergetic needs. For example, if media with relatively low glucose and/or glutamine were replaced at a certain interval, the glucose and/or glutamine could repeatedly reach restricted levels before being replaced without total cell death occurring. We sought to test both conditions.

First, we conducted an 8-day media change experiment in the CT-2A and the VM-M3 cells cultured either in low glucose, or in low glucose with the addition of various concentrations of lactate, both under physiological glutamine levels (Figure 4A and 4B). Media was changed every 48 hours. For both the CT-2A and VM-M3 cells, bioluminescence in lactate-containing media was indistinguishable from bioluminescence in identical media without lactate, regardless of lactate concentration. Bioluminescence was low (but non-negligible) in all low glucose conditions, regardless of lactate supplementation, suggesting partial loss of cell viability from glucose deprivation. Next, we restricted glucose and glutamine to varying degrees in U-87MG (Figure 4C). The addition of lactate had no effect on U-87MG cell bioluminescence in media with 5.0 mM glucose and 1.0 mM glutamine. However, under severe nutrient restriction (0.5 mM glucose and 0.5 mM glutamine), the addition of 5 mM lactate resulted in a 46% increase in bioluminescence, with higher concentrations of lactate failing to induce further increases.

**Figure 4.**
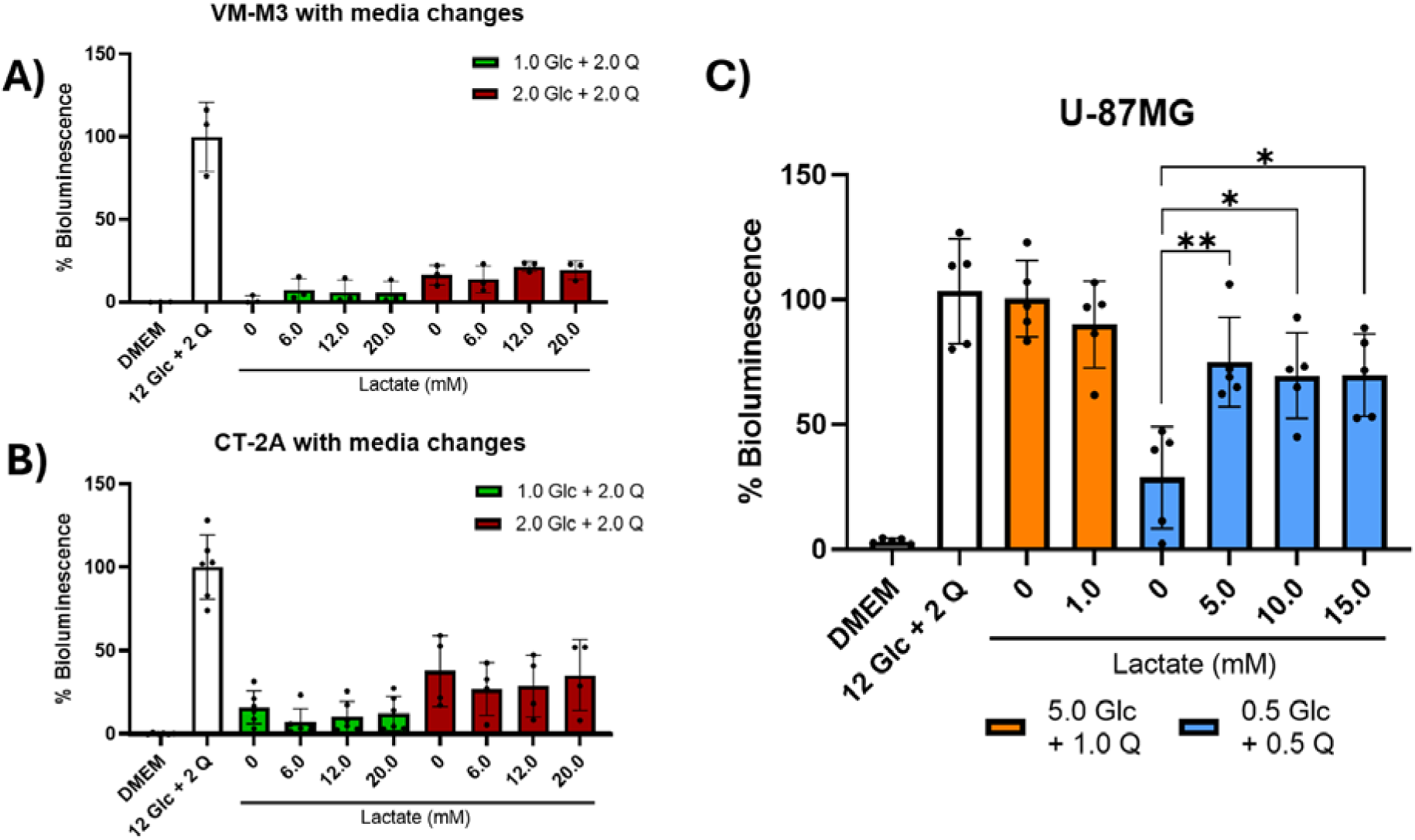
Influence of lactate on ATP content and viability with intermittent media replacement and restriction of glucose in mouse and human glioma cells. (A) VM-M3 glioma cells were seeded and cultured for 8 days in the specified media. Media was replaced every 48 hours. (B) CT-2A glioma cells were seeded and cultured for 8 days in the specified media. Media was replaced every 48 hours. (C) U-87MG cells were seeded and cultured for 72 hours in the specified media. All values are expressed in millimolar (mM). All quantifications were normalized to the positive control (12.0 mM glucose and 2.0 mM glutamine). Error bars represent mean ± SD with a minimum of 3 independent experiments. Significance levels are denoted as ∗ *p* ≤ 0.05 and ∗∗ *p* ≤ 0.01. The results show that lactate had no significant effect on ATP content and viability in the VM-M3 or CT-2A cells regardless of glucose restriction, while lactate increased ATP content and viability in the U-87MG cells only under conditions of severe glucose restriction.

### 2-deoxy-D-glucose eliminates the effects of lactate on viability and oxygen consumption rate in U-87MG glioma cells

Given the effect of lactate on U-87MG viability under restricted glucose, we aimed to assess whether this effect would remain if the glycolysis pathway itself was simultaneously inhibited. We used 2-deoxy-D-glucose (2DG), a well-characterized glycolysis inhibitor [54]. We selected 0.25 mM 2DG after performing a dose-response curve to calculate the IC50 of 2DG in 0.5 mM glucose and 0.5 mM glutamine at 72 hours (Figure S6). As this is a low concentration of 2DG relative to the 1.0 to 10.0 mM typically used in *in vitro* cancer research [55], we also selected two higher concentrations of 2DG: 2.0 mM and 5.0 mM. As above, with a baseline medium containing restricted glucose, U-87MG cells cultured in an additional 15.0 mM of lactate exhibited a significant increase in bioluminescence compared to cells without lactate (Figure 5A). However, this effect was abolished by the addition of as little as 0.25 mM 2DG. At the same time, bioluminescence was similar in the U-87MG cells cultured in the presence (0.25 mM) or absence of 2DG in the restricted baseline media. These findings showed that a low concentration of 2DG could abolish the effect of lactate on U-87MG viability without significant toxicity.

**Figure 5.**
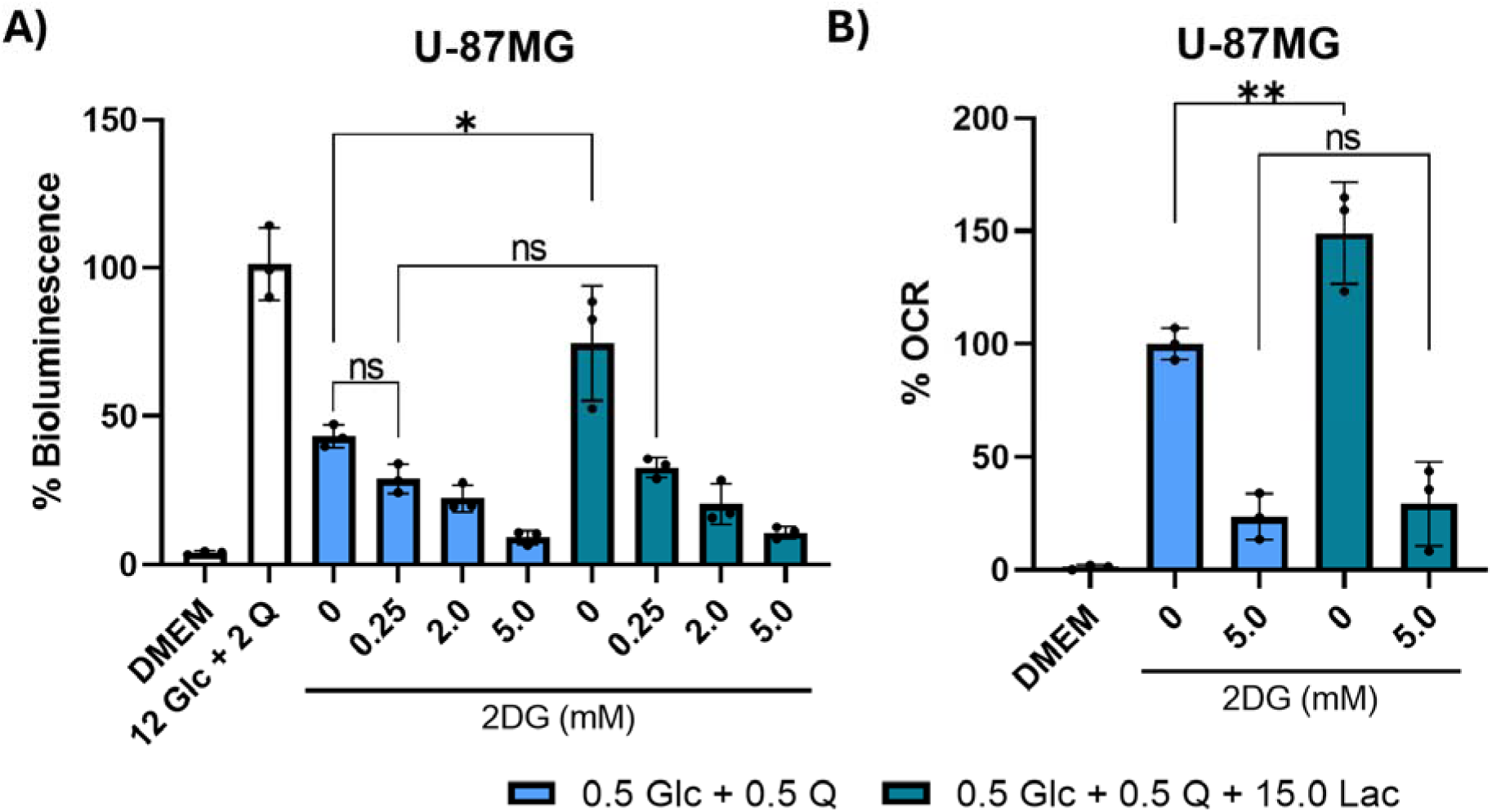
Influence of lactate and 2-deoxy-D-glucose on ATP content, viability, and oxygen consumption rate in U-87MG glioma cells. (A) U-87MG cells were seeded and cultured for 72 hours in the specified media. Bioluminescence was normalized to the positive control (12.0 mM glucose and 2.0 mM glutamine). (B) U-87MG cells were seeded and then cultured for 72 hours while measuring OCR using the Resipher instrument. OCR was normalized to cells cultured in 0.5 mM glucose and 0.5 mM glutamine. All values are expressed in millimolar (mM). Error bars represent mean ± SD with a minimum of 3 independent experiments. Significance levels are denoted as ∗ *p* ≤ 0.05 and ∗∗ *p* ≤ 0.01. The results show that 0.25 mM of 2DG abolished the effect of lactate on U-87MG ATP content and viability with minimal toxicity, while 5 mM of 2DG abolished the effect of lactate on OCR.

While it should be noted that oxygen consumption rate (OCR) does not necessarily indicate ATP synthesis via OxPhos [5], any OxPhos that does occur should be accompanied by an increase in OCR. Lactate is non-fermentable in mammals and must be oxidized to contribute to ATP synthesis as a metabolic fuel. Thus, if U-87MG utilized lactate as a fuel, this should be accompanied by a relative increase in OCR due to increased OxPhos. We measured OCR with the addition of lactate and 2DG to determine whether lactate could increase OCR and whether 2DG could abolish this effect (Figure 5B). Cells in the restricted baseline media with added lactate demonstrated a 49% increase in OCR compared to cells in the restricted media without lactate. This effect was abolished upon 2DG treatment, suggesting that any lactate-dependent OCR was ultimately dependent on glycolysis.

## Discussion

Our results showed that lactate was unable to serve as a major metabolic fuel or substitute for glucose to support viability in either the mouse (VM-M3 and CT-2A) or the human (U-87MG) glioma cells. Lactate alone failed to sustain viability of these glioma cells at any concentration. Lactate as a supplement to glutamine also failed to increase viability except with the U-87MG cells cultured under severe glucose restriction. The latter observation in U-87MG cells might be due in part to (a) the 7-fold slower basal metabolic rate in humans than in mice [56,57], or (b) the relatively high expression of lactate dehydrogenase B (LDHB) in U-87MG [57,58]. Moreover, the lactate effect observed in the human U-87MG cells was abolished by a low 0.25 mM dose of the glycolysis inhibitor 2DG, suggesting that any lactate utilization was dependent on glycolysis. Notably, cell culture dosages of 2DG typically range from 1.0 mM to 10.0 mM [55], but the selected 0.25 mM dose of 2DG could have been potentiated by lower competition for glucose transporters and glycolytic enzymes due to glucose deprivation [54]. A failure to entirely remove glucose and glutamine from culture media together with an assumption that OCR is an accurate marker for ATP production through OxPhos might have contributed to previous suggestions that lactate could fuel cancer growth [24–26]. Recent information shows that the fermentation of glucose and glutamine together are necessary and sufficient for tumor cell growth, whereas ATP production via OxPhos is neither necessary nor sufficient [13,52]. Our findings are consistent with this concept as we found that lactate was not a major oxidizable fuel for maintaining glioma viability.

In contrast to the mouse and human glioma cells, we found that lactate could significantly enhance viability in the non-neoplastic VM/Dk and C57BL/6J TPMs. Notably, the C57BL/6J and the VM/Dk TPMs cultured in glutamine and lactate reached viability levels comparable to TPMs cultured in glucose and glutamine, indicating that lactate could replace glucose to support viability in these non-neoplastic TPMs. These differences between the neoplastic and non-neoplastic cells are unlikely to result from genetic variation, as the C57BL/6J and the VM/Dk TPMs were syngeneic with the mouse CT-2A and VM-M3 gliomas, respectively. Due to their intact OxPhos, the TPMs would be expected to oxidize lactate more efficiently than the glioma cells. Indeed, our findings are consistent with previous research showing that bone marrow-derived macrophages could oxidize lactate *in vitro* for ATP production [59]. Hence, our findings take on added significance as we compared neoplastic tumor cells with non-neoplastic syngeneic control cells [60].

Although our findings failed to support a major role for lactate as a fuel for ATP production and viability in the glioma cell lines that we evaluated, our findings do not exclude other roles for lactate in cancer. Torrini et al. found increased bioluminescence in U-87MG cells cultured in 0.5 mM glucose and 0.5 mM glutamine when supplemented with at least 10 mM lactate [25]. Cai et al. found that HepG2 cells survived and proliferated for 8 days when cultured under relative glucose restriction with 20 mM L-lactate supplementation and media replacement every 48 hours [24]. Cai et al. also found that D-lactate could increase HepG2 proliferation, reduce extracellular acidification, and increase OCR [24]. As D-lactate is not metabolized by mammalian LDHA or LDHB [61], these findings suggest that lactate may have signaling effects independent of OxPhos [24]. As our study focused on physiologically relevant conditions, we used only the L-lactate isomer. Likewise, both the Cori Cycle and TME acidification are critical for *in vivo* lactate metabolism [35,62–64]. Our *in vitro* system with L-lactate supplementation excluded both as potential explanations of our findings. Lactate and succinate are the end products of the high-throughput glycolysis and glutaminolysis pathways, respectively, which drive the dysregulated growth of cancer cells [13,52]. It is well known that the tumor secretion of lactate (with a proton) together with succinate will acidify the TME, thus contributing to cancer progression [63]. The lactate produced in cancer cells can also return to the tumor as glucose via the Cori cycle, thus maintaining a constant supply of glucose to the tumor [35,62–65]. Reports of L-lactate contributing to cancer progression *in vivo* may be attributable to signaling pathways, the Cori cycle, or TME acidification, rather than lactate oxidation. Our findings do not support a heterocoprophagic mechanism where lactate detritus produced in normal fibroblasts would fuel lactate-producing tumor cells through OxPhos [26]. An increase in OCR following lactate supplementation also does not necessarily indicate efficient ATP synthesis via complex V, as OCR can be uncoupled from OxPhos, such as when oxygen is diverted to produce reactive oxygen species or during proton gradient uncoupling [5]. While labeled lactate carbons can enter the TCA cycle, the extent of this contribution remains controversial and does not indicate ATP synthesis via OxPhos alone is sufficient to maintain proliferation [22,23,37,38,66]. Thus, while lactate may have several indirect roles in supporting cancer cell metabolism, our findings do not support lactate as a major oxidative fuel in glioma. We conclude that therapeutic efforts should prioritize glycolysis and glutaminolysis targeting over lactate oxidation.

A limitation of our study was that we evaluated only three glioma cell lines, with U-87MG as the sole human-derived line. However, the metabolic abnormalities found across these models are similar to those found in the cells of all major cancers [13,52]. Nevertheless, it would be important to determine if our findings could be replicated in cell lines from other cancers.

## Conclusion

Our data indicate that lactate does not serve as a significant metabolic fuel for neoplastic glioma cells. Lactate alone failed to sustain viability across all tested mouse and human glioma cell lines. Lactate was unable to substitute for glucose to support viability in the presence of glutamine. The addition of lactate to glutamine had no effect on viability, except under severe glucose restriction in U-87MG cells. In this case, lactate increased viability and oxygen consumption rate, but both effects were abolished by a low dose of 2DG, suggesting that the contribution of lactate was glycolysis-dependent. In contrast to the glioma cells, lactate significantly enhanced viability in primary TPMs, suggesting fundamental metabolic differences between neoplastic and non-neoplastic cells. Our findings support the perspective that glucose and glutamine are the predominant drivers of ATP production in glioma cells. While our results do not exclude non-oxidative roles for lactate in glioma progression, they suggest that targeting glycolysis (either via nutrient depletion or pharmacological inhibition) should take precedence over targeting lactate oxidation when designing rational metabolism-based therapeutic approaches.

## Acknowledgments

We thank the Foundation for Metabolic Cancer Therapies, The Nelson and Claudia Peltz Family Foundation, Lewis Topper, The John and Kathy Garcia Foundation, Dr. Edward Miller, Kenneth Rainin Foundation, the Corkin Family Foundation, Children with Cancer UK, the Broken Science Initiative, Delaware County Special Deputies Benevolent Fund, and the Boston College Research Expense Fund for their support. The funders had no role in study design, data collection and analysis, decision to publish, or preparation of the manuscript.

## Author contributions

EYM, TD, NLT, DCL, PM, OS, and TNS contributed to the conceptualization and design of the research. EM performed the bulk of experiments and acquired the data. EYM, TD, NLT, DCL, and TNS drafted and revised the manuscript. All authors have accepted responsibility for the entire content of this manuscript and approved its submission.

## Data availability

The raw data can be obtained on request from the corresponding author.

## Abbreviations

EthD-III: Ethidium homodimer III
OxPhos: Oxidative phosphorylation
TPMs: Thioglycollate-elicited peritoneal macrophages
OCR: Oxygen consumption rate
2DG: 2-deoxy-D-glucose
LDH: Lactate dehydrogenase
GBM: Glioblastoma
TME: Tumor microenvironment
DMEM: Dulbecco’s Modified Eagle Medium
Lac: Lactate
Glc: Glucose
Q: Glutamine

