## Supplementary Figures for "Lactate cannot replace glucose for maintaining the viability of mouse and human glioma cells"

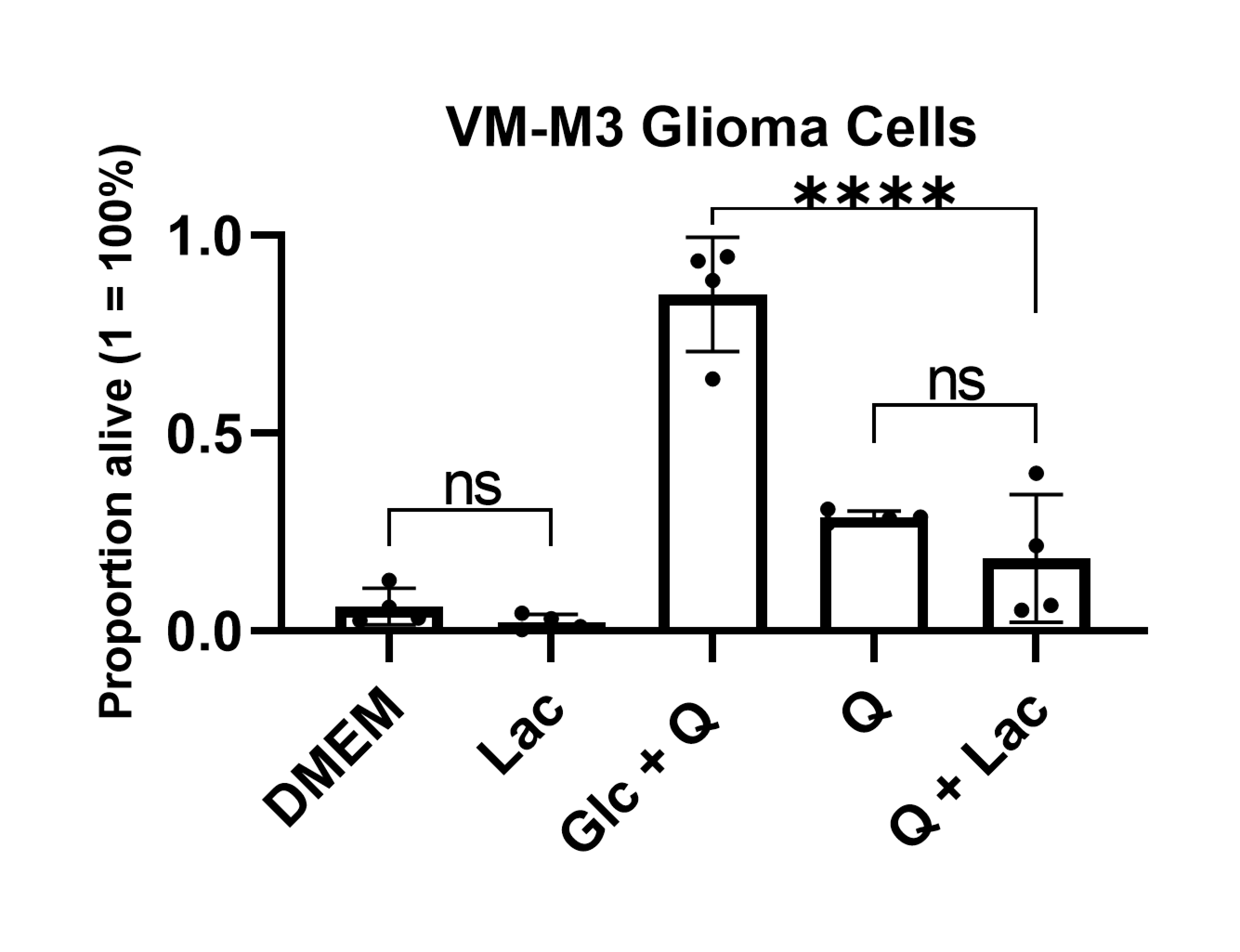

**Figure S1**. Validation of bioluminescence as a proxy for viability of VM-M3 glioma cells via quantification of calcein-AM and EthD-III fluorescent double-staining assay. VM-M3 cells were seeded and cultured for 48 hours in the specified media. Viability was quantified via CellProfiler using EthD-III and calcein-AM staining images. The media conditions were DMEM (negative control), glucose and glutamine (positive control; Glc + Q), lactate (Lac), glutamine (Q), or glutamine and lactate (Q + Lac). When present, glucose, lactate, and glutamine concentrations were 12 mM, 12 mM, and 2 mM, respectively. Error bars represent mean ± SD with a minimum of 4 independent experiments. These results align with the bioluminescence results for the VM-M3 glioma cells and demonstrate the same patterns of significance across the relevant comparisons, indicating that ATP-linked bioluminescence is a proxy for viability.

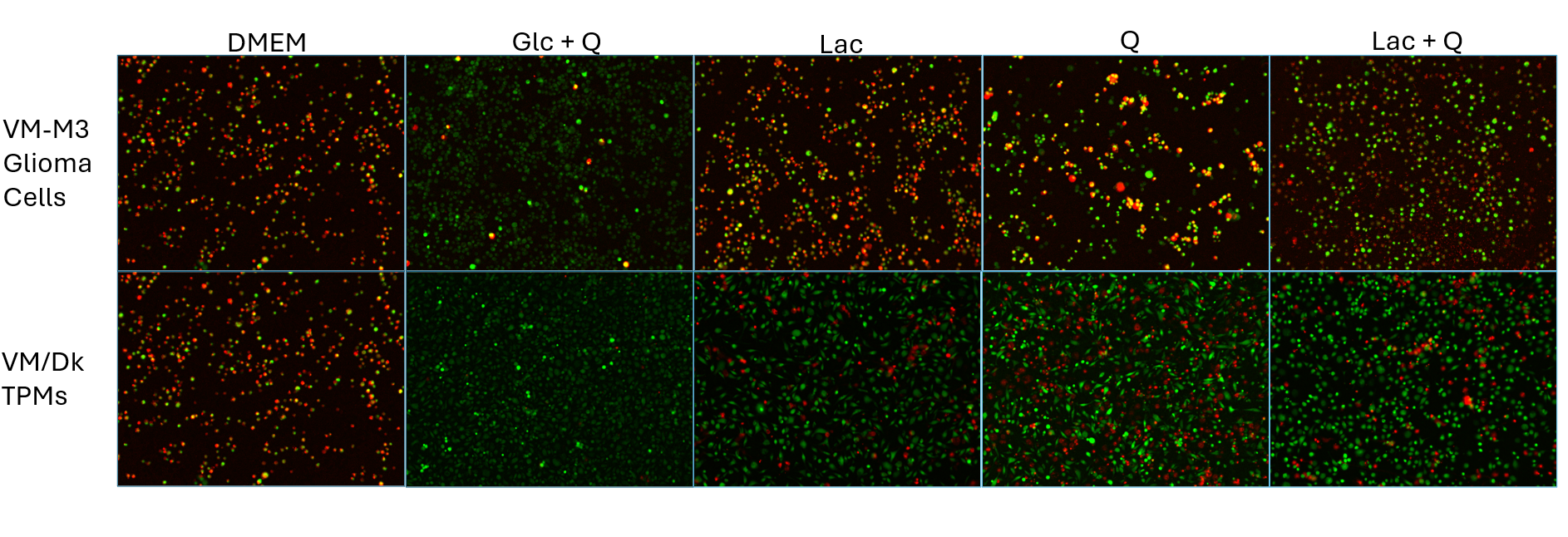

**Figure S2**. Representative calcein-AM and EthD-III fluorescent double-staining images of the VM-M3 glioma cells and VM/Dk TPMs. EthD-III only enters cells when the plasma membrane is compromised and fluoresces red when it binds to nucleic acids, thus marking dead or late apoptotic cells. Calcein-AM is a membrane-permeable dye that is hydrolyzed by intracellular esterases in live cells to produce green fluorescence. Therefore, cells exhibiting only red fluorescence are non-viable and cells emitting only green fluorescence are viable (see Methods). These results illustrate the contrast in viability between the VM-M3 glioma cells and the VM/Dk TPMs in lactate-containing media, showing near-complete cell death of VM-M3 cells cultured in lactate alone, whereas 78% of VM/Dk TPMs survived in lactate alone.

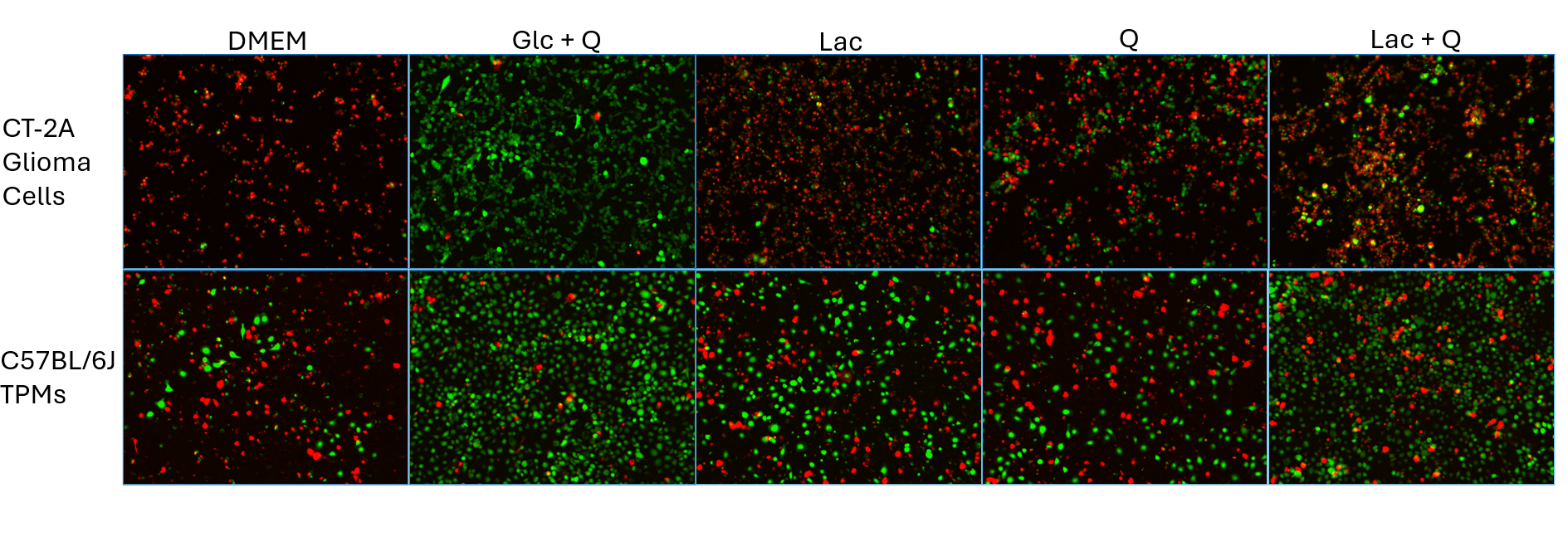

**Figure S3**. Representative calcein-AM and EthD-III fluorescent double-staining images of the CT-2A glioma cells and C57BL/6J TPMs. EthD-III only enters cells when the plasma membrane is compromised and fluoresces red when it binds to nucleic acids, thus marking dead or late apoptotic cells. Calcein-AM is a membrane-permeable dye that is hydrolyzed by intracellular esterases in live cells to produce green fluorescence. Therefore, cells exhibiting only red fluorescence are non-viable, cells emitting only green fluorescence are viable (see Methods). These results illustrate the contrast in viability between the CT-2A glioma cells and the C57BL/6J TPMs, showing near-complete cell death of CT-2A cells cultured in lactate alone, whereas 48% of C57BL/6J TPMs survived in lactate alone.

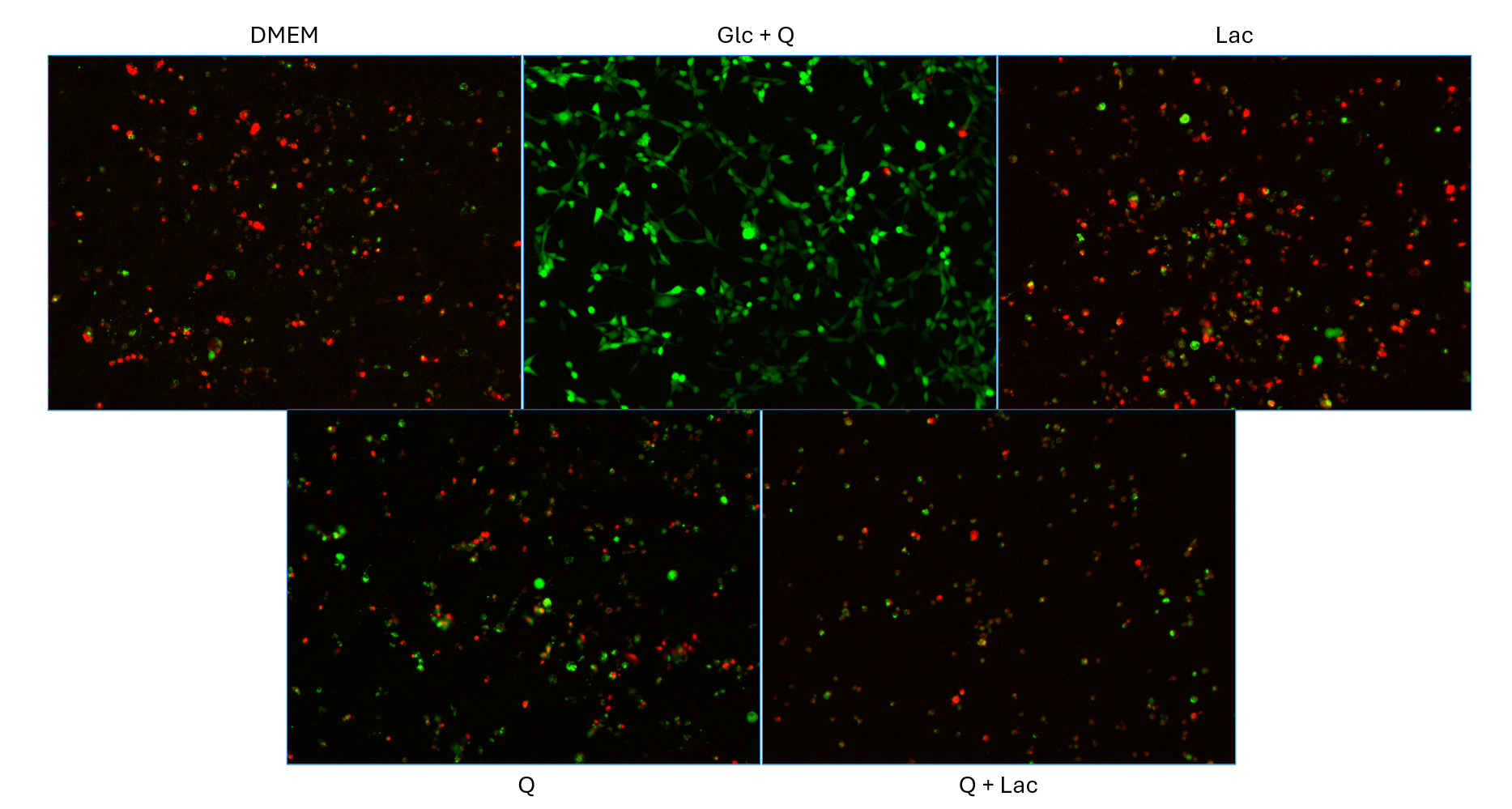

**Figure S4**. Representative calcein-AM and EthD-III fluorescent double-staining images of the U-87MG glioma cells. EthD-III only enters cells when the plasma membrane is compromised and fluoresces red when it binds to nucleic acids, thus marking dead or late apoptotic cells. Calcein-AM is a membrane-permeable dye that is hydrolyzed by intracellular esterases in live cells to produce green fluorescence. Therefore, cells exhibiting only red fluorescence are non-viable, cells emitting only green fluorescence are viable (see Methods). These results illustrate the near-complete cell death among U-87MG cells cultured in lactate alone and glutamine with lactate.

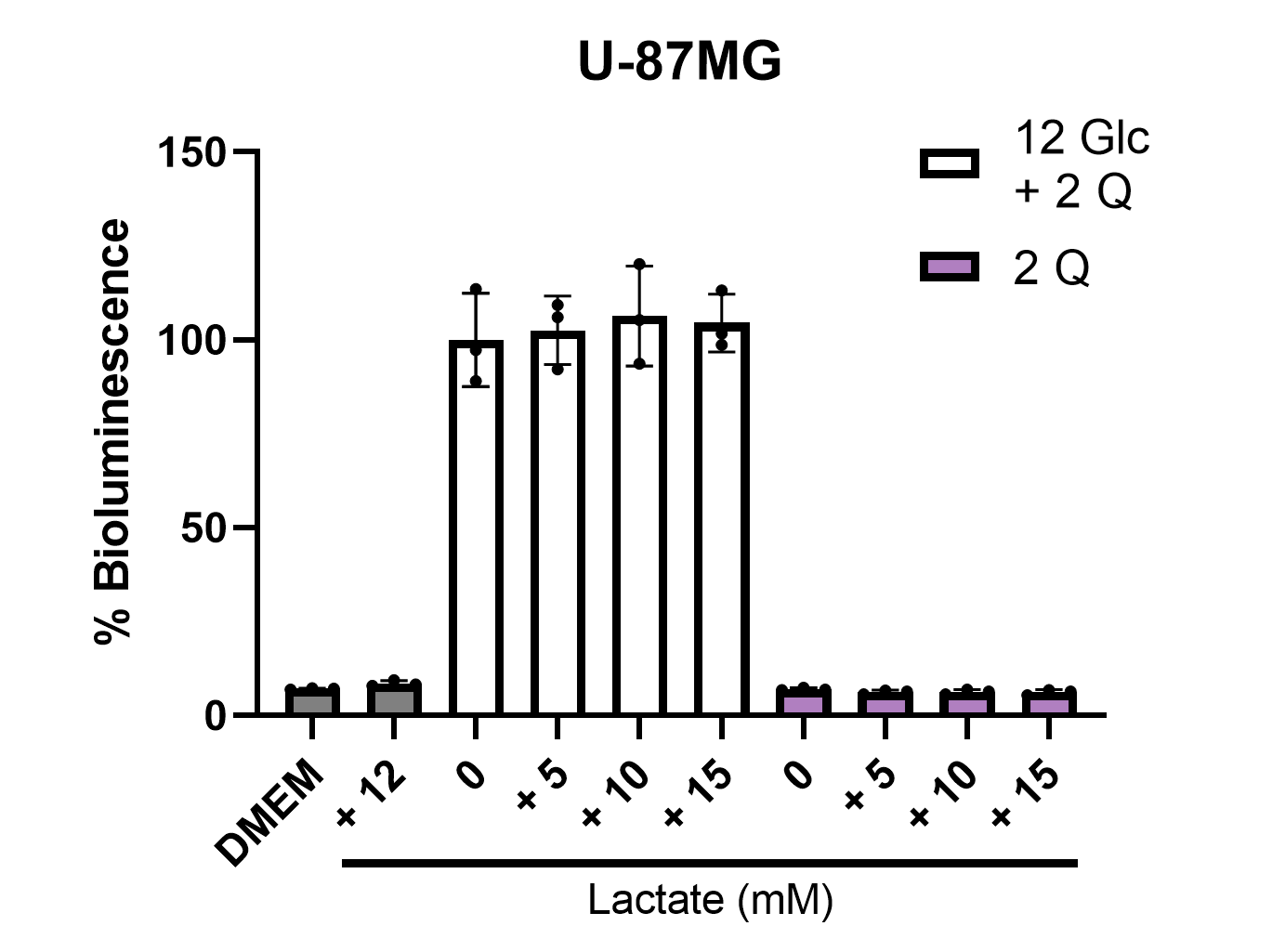

**Figure S5**. Effects of different concentrations of supplementary lactate on ATP content and viability in U-87MG. U-87MG glioma cells were seeded and cultured for 72 hours in the specified media. When present, glucose and glutamine concentrations were 12 mM and 2 mM, respectively. All quantifications were normalized to the positive control (12 mM glucose and 2 mM glutamine). All values are expressed in millimolar (mM). Error bars represent the mean ± SD from at least three independent experiments. There was no effect with the addition of lactate in any case, regardless of concentration.

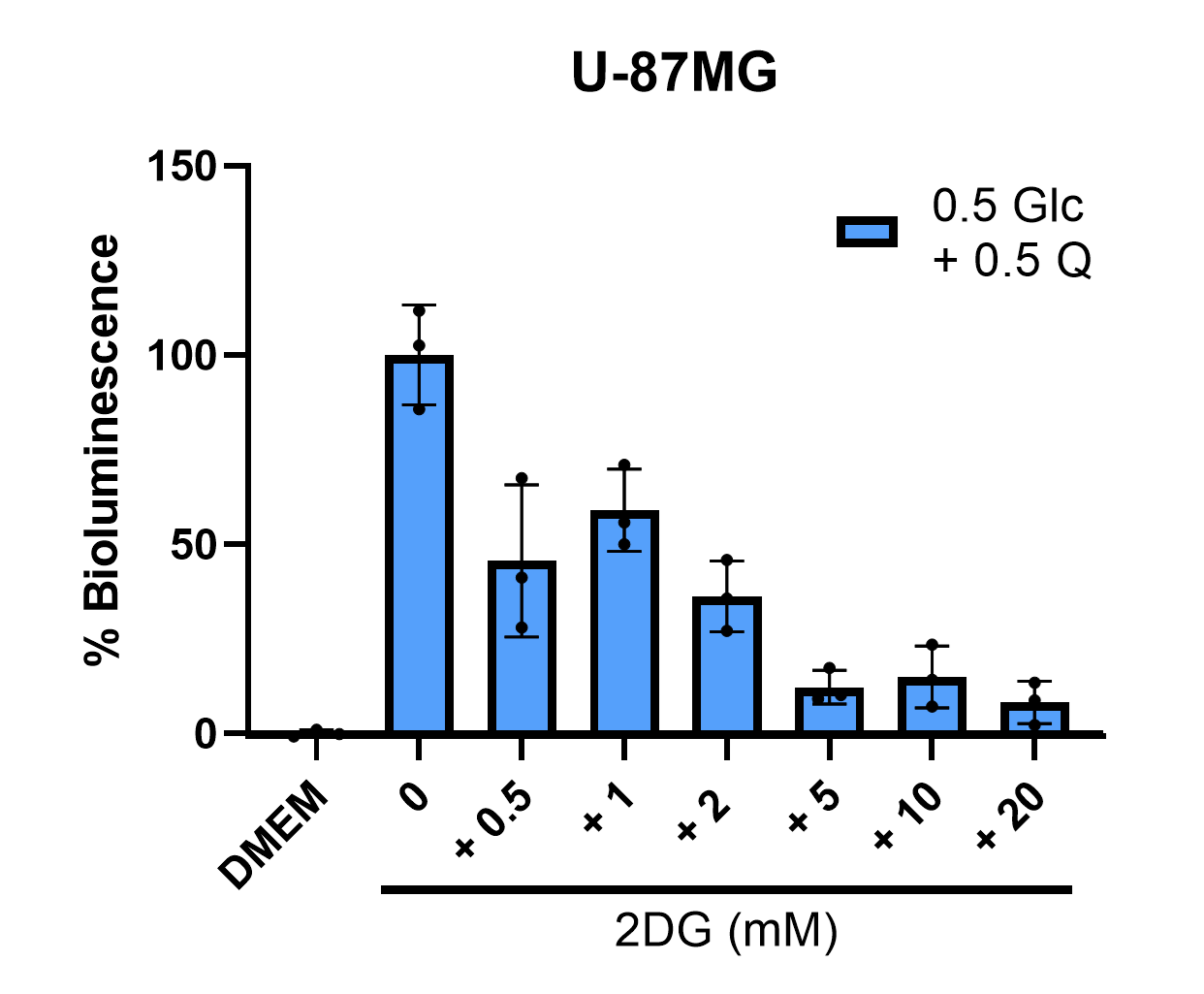

**Figure S6**. Dose-response curve of 2DG in 0.5 mM glucose and 0.5 mM glutamine. U-87MG glioma cells were seeded and cultured for 72 hours in the specified media. This dose-response curve was used to calculate the IC50 of 2DG in 0.5 mM glucose and 0.5 mM glutamine, which was 0.2866 mM. All quantifications were normalized to 0.5 mM glucose and 0.5 mM glutamine. All values are expressed in millimolar (mM). Error bars represent the mean ± SD from at least three independent experiments.
